# Furin-cleavage site is present in an antiparallel β-strand in SARS-CoV2 Spike protein

**DOI:** 10.1101/2022.02.18.481028

**Authors:** Arif Bashir, Naveed Nazir Shah

## Abstract

Furin cleavage-site (CS) present between the S1/S2 junction in SARS-CoV2 spike (S) protein is critical to drive the fusion of SARS-CoV2 with the host cell. SARS-CoV2 falls in the sarbecovirus lineage that doesn’t comprise of furin CS and therefore makes its origin enigmatic. The available wild-type (Wt) SARS-CoV2 S protein with PDB ID: 6yvb lacks a stretch of amino acid including furin CS as well. All investigators till date have shown this stretch existing in the form of a loop. We are for the first time reporting that this stretch comprises of 14 amino acid residues (677QTNSPRRARSVASQ689), forming an antiparallel β-sheet comprising of PRRAR furin CS. We observed the presence of this antiparallel β-sheet in MERS spike protein as well. While switching over from Wt. SARS-CoV2 with PRRAR furin CS to B.1.1.7 variant with HRRAR furin CS, we found 3% increase in the percentage content of β stands. Interestingly, we found that the change of B.1.1.7 to B.1.617 variant comprising of RRRAR furin CS shifted the percentage secondary structure back to that found in Wt. SARS-CoV2. We anticipate that this β-sheet is used as a docking site by host cell proteases to act on furin-CS. Additionally, we studied the interaction of modeled SARS-CoV2 S protein with transmembrane protease, serine 2 (TMPRSS2), and furin proteases, which clearly highlighted that these proteases exclusively uses furin CS located in β-sheet to cleave the SARS-CoV2 S protein at its S1/S2 junction.

## Introduction

### Severe acute respiratory syndrome coronavirus-2 (SARS-CoV-2)

Severe acute respiratory syndrome coronavirus-2 (SARS-CoV-2) caused a local outbreak in Wuhan city in December 2019, which subsequently, at an alarming rate, gripped the other continents, pressing the World Health Organization to declare it a pandemic [1-3]. This virus belongs to a betacoronavirus genus that includes SARS-CoV, Middle-East Respiratory Syndrome (MERS-CoV), and bat SARS-related CoVs. SARS-CoV-2 RNA genome sequence shares ∼80% sequence similarity with the SARS-CoV, and ∼50% with the MERS-CoV. Interestingly, the bat CoV RaTG13 is the only nearby relative of SARS-CoV2 that shares 93.1% sequence similarity with the spike (S) protein compared to other CoVs [4]. SARS-CoV2 S protein is composed of 1092 amino acid and is divided into subunit 1 (S1) and S2 region. SARS-CoV2 S protein has accumulated maximum number of mutation during the last two years. Wild-type SARS-CoV2 that emerged in late 2019, right now has more than eight variant of concern identified in different countries namely, B.1.1.7 (UK-Variant), B.1.1.318. (UK-Variant), B.1.351/501.V2 (South-African Variant), B.1.429 (United-States Variant), B.1.617 (Double-mutant; Indian-Variant), B.1.618 (Triple-mutant; Indian-Variant), B.1.617.2 (Delta variant), Delta plus (AY.1), and B.1.525 (UK-Denmark Variant), and P.1 (Brazilian Variant) [5]. The emergence of VOC from the original wild-type SARS-CoV2 is clearly reflecting towards the selective pressure build on it that consequently gave rise to quasispecies viral population [6, 7]. Use of Remidesivir, Interferon (IFN) α, IFNβ, IFNγ, convalescent plasma, and persistent infection in the host are some of the driving forces behind the emergence of quasispecies viral population outcompeting the dominant wild-type SARS-CoV2 are resistant to many therapies, which is really a matter of concern and should be taken into account while putting on the infected patients on a number of antiviral drugs, antibiotics or cocktail of them [7].

### SARS-CoV2 Spike Glycoprotein

SARS-CoV2 entry into host cells is facilitated by spike glycoprotein present on its envelope. This protein forms a homotrimer structure projecting out from its main body. This projected spike protein is an attractive target to generate antibodies and antiviral agents that will hampers it association with the host cell [8]. The SARS-CoV2 spike (S) protein comprises of two functional subunits, namely, S1 and S2 subunits. The S1 subunit comprises of N-terminal domain and the receptor binding domain (RBD) [9]. The S1 subunit of SARS-CoV2 first binds with the ACE receptor present on the host cell. The S2 subunit is composed of many regions that include fusion peptide (FP), heptad repeat 1, central helix, connector domain, heptad repeat 2, transmembrane domain, and cytoplasmic tail [10] (**Fig 1A**). This subunit finally promotes the fusion the virus with the host cells. Coronaviruses get access into the host cell through their spikes on their surface. Ordinarily, S protein of coronaviruses is inactive prior to its binding with the host cell. Upon binding with the host cell receptor, coronaviruses achieve the ability to mediate their fusion with the host cell membrane by cleavage at S1/S2 cleavage site (CS) present in the S protein. Depending upon the uniqueness of the respective amino acid sequence at S1/S2 CS in the S protein of coronaviruses, the cleavage is achieved by: (1) transmembrane serine-proteases (2) furin-like enzymes in the host cell (3) cathepsin proteases in the late endolysosome/endosome. Post S1/S2 site cleavage, a second cleavage site in the S2 domain of the spike protein gets exposed to serine-proteases or cathepsins, which subsequently drives the host-virus membrane fusion [11]. Cryo-electron microscopy performed on the SARS-CoV S protein bound with the ACE2 has revealed that initially S1 binds with the ACE2 receptor on the host cell receptor. This binding then drives the disassociation of S1 with the ACE2 receptor that consequently prompts the S2 to shift from a metastable prefusion to a more stable post-fusion state that drives the membrane fusion [8, 9, 12, 13]. In the prefusion conformational state, S1 and S2 subunits remain non-covalently bound. The S1 subunit comprising of the RBD houses receptor binding motif (RBM) (438−506) that interacts with the N-terminal peptidase domain of ACE receptor (**Fig 1B**). SARS-CoV2 S protein exists in closed and the open conformational states. SARS-CoV2’s RBD has higher binding affinity with the ACE2 receptor compared to SARS-CoV. SARS-CoV-2’s F486 interacts with the Q24, L79, M82, and Y83 amino acids of the ACE2 receptor compared to SARS-CoV’s L472 that interacts with the L79 and M82 amino acids of the ACE2 receptor (**Fig 1C**). The Q493 amino acid of the SARS-CoV2 RBD interacts with the ACE2’s K31, E35, and H34 amino acids. In RBM, ACE’s D30 forms a salt-bridge with the SARS-CoV-2 K417. Genetically engineered SARS-CoV2 RBD−ACE receptor complex comprising of RBM and the RBD core of the SARS-CoV S protein acts as a scaffold to promote crystallization. The side loop pointing away from the main binding region preserves a salt-bridge (N-O) between SARS-CoV-2 RBD’s R426 and E329 of ACE receptor. The claw-like exposed structure of the ACE receptor binds with the concave SARS-CoV-2 RBM. The binding affinity of the ACE2 receptor with the SARS-CoV RBD is less compared to the SARS-CoV-2 and therefore, the infectivity and pathogenesis of SARS-CoV is less than the SARS-CoV2 however both recognizes the seventeen amino acid residues present in the human ACE2 receptor. In the closed state, the three RBM do not project out and all the three RBDs are facing down. In the open state, one of the RBD is in the upward conformation, which is necessary for the SARS-CoV-2 to fuse with the host cell [10] (**Fig 1D, E**).

**Fig 1.**
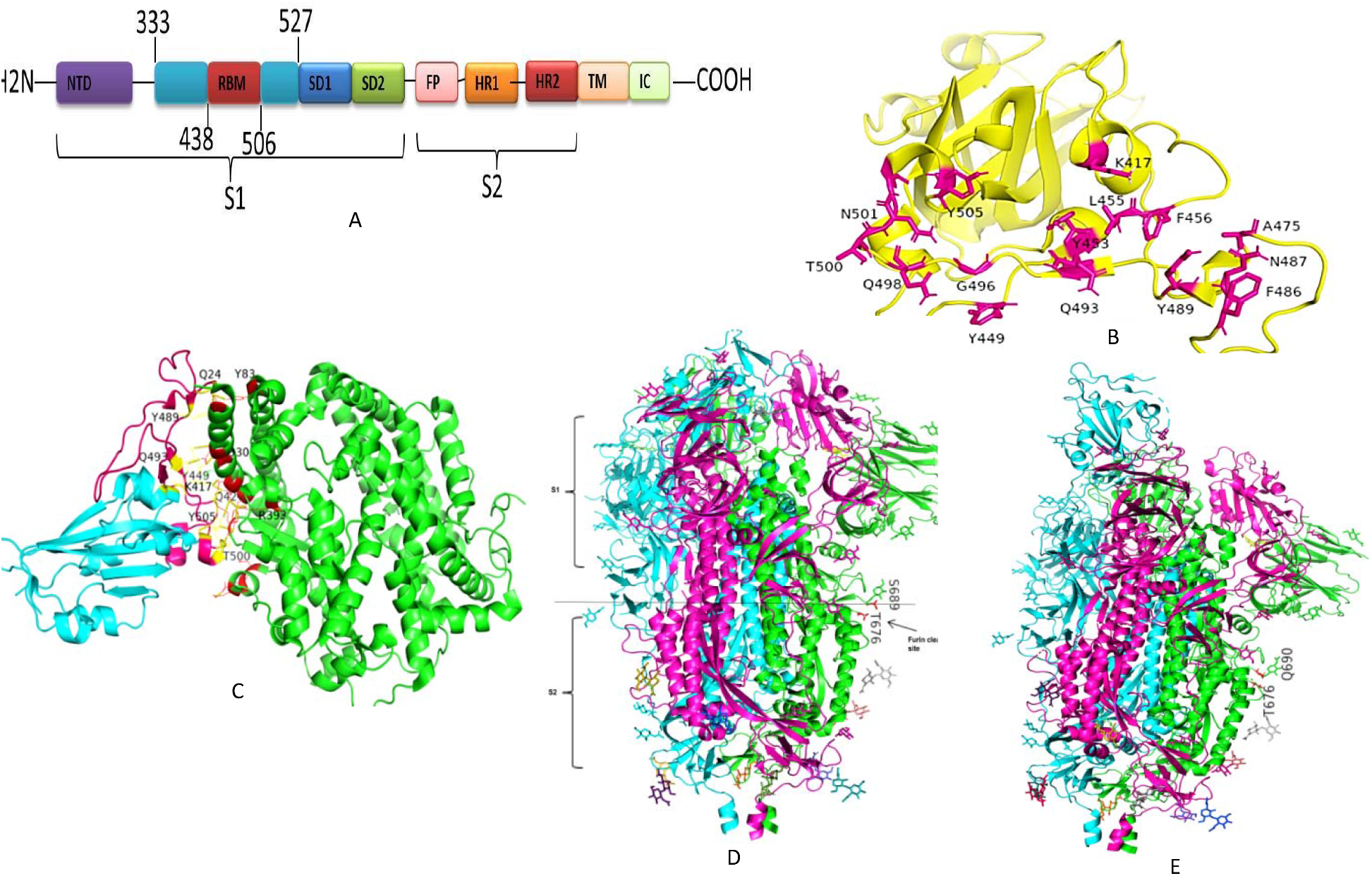
(**A**) Structural organisation of SARS−CoV2 spike monomer. FP: Fusion peptide, HR1: Heptad repeat1, HR2: Heptad repeat2, IC: Intracellular domain, NTD: N-terminal domain, SD1: Subdomain 1, SD2: Subdomain 2, and TM: Transmembrane region (**B**) Critical amino acids (Highlighted with magenta color) present in the SARS−CoV2 RBD (**PDB: ID 6M0J**) important for its interaction with ACE2 receptor. (**C**) Interaction between SARS-CoV2-RBM (receptor binding motif, highlighted with magenta color) with the critical amino acids of ACE2 receptor (highlighted with red color; PBD: 6M0J). **(D)** SARS-CoV2 trimeric spike protein closed (**PDB: 6VXX**) and (**E**) open (**PDB**: **6VYB**) conformation.

The secondary structure of the RBD consists of five β-strands organized in antiparallel fashion that is, in turn, connected by dominating loops (stabilized by disulphide linkage) and short α-helices. Among the five β-strands, β4 and β7 consists of RBM, which in turn consists of α-helices, loops, and a short β5 and β6 strands. RBM is the portion of RBD that comprises of most of the binding sites for the human ACE2 receptor peptidase domain (19− 615)[10]. The SARS-CoV-2 S protein has nine cysteine residues and among them eight forms the four pair of disulphide linkage. The three disulphide linkage (C336-C361, C379-C432, and C391-C525) in the RBD increases the stability of β sheet present in it while as the C480-C488 linkage assists in promoting the connection between the loops present in the RBM. The crystallographic data has revealed the homotrimeric nature of the SARS-CoV2 S protein and among them one S protein’s RBD is in up conformation or in open/ active state[14]. The binding of open conformation of the S protein with S1 subunit destabilizes the prefusion structure. Subsequently the S1 subunit detaches from the ACE2 receptor−S protein that promotes the S protein S2 subunit to refold into a stable post-fusion state. The SARS-CoV2 RBD straddles between a hinge-like structure that exposes or hides the RBD’s active site to interact or disengage with the ACE receptor at the lung epithelial cells[10, 15].

### SARS-CoV-2 furin cleavage site

One of the remarkable features of the SARS-CoV-2 S protein is the presence of a four-amino acid sequence that starts with an amino acid acid proline (SPRRAR|S) that lies between the RBD (S1) and fusion (S2) domain [16]. This amino acid sequence is referred as a polybasic site and has been anticipated to be a part of the closely related origin of the SARS-CoV2 pandemic around the world[17]. Proteolytic cleavage of the S protein at its S1/S2 and S2 junction is extensively used to trigger the fusion apparatus of viral glycoproteins [11, 18]. SARS-CoV-2 S protein cleavage is not a simple process rather it involves a series of complex process comprising of more than one cleavage events at different sites coupled with the participation of wide range of host cell proteases [19]. Furin, a host cell protease, is ubiquitously expressed at low levels in Golgi apparatus of all cells. It plays an important role in virus-driven pathogenesis with its ability to cleave polybasic amino acid cleavage site (CS) found in some avian influenza virus and generally present in betacoronaviruses, thereby allowing its systemic dissemination. These polybasic sites are typically generated through polymerase slippage during replication of some avian influenza virus. Few viruses, such as influenza virus, are devoid of polybasic sites, and infection remains localized to those cells that possess trypsin-like proteases. Bat-origin MERS-like merbecovirises also harbors furin CS. Interestingly, it has been reported that the simple insertion of a polybasic site in H3 virus doesn’t led to a high degree of pathogenicity and has been likely originated through a series of genetic changes during the course of a natural selection. However, as far as the origin and pathogenicity of SARS-CoV2 through recombination/mutation of a bat-origin virus has been understand as obscure [20]. The acquisition of furin cleavage motif between S1-S2 junction of SARS CoV2 S protein is astonishing and has led us to speculate that this virus has been genetically engineered. The furin cleavage sequence in SARS-CoV2 S protein doesn’t resemble with the prototypical furin CS RxK/RR found in H3 virus. The wild-type (Wt) SARS-CoV2 that first emerged had proline residue in furin CS. Subsequently, this residue was replaced by histidine in B.1.1.7 variant, and arginine in B.1.617 where tri-basic PRRAR/S sequence to RRRAR/S along with other adaptions. One can fairly infer that progressively these sites are becoming more polybasic with the continuous pandemic and increasing transmissibility that may consequently leads to the development of new variants [18].

Progressive accumulation of mutation in the spike protein has added a layer complexity and has raised many questions such as how the S protein adapts to the environment of different species, tissue-types, and cell types. In the middle of the SARS-CoV2 S protein comprises of furin cleavage site for the subtilisin-identical host cell protease furin. Ordinarily host proteases, more particularly, TMPRSS cleaves the spike glycoprotein at the S2 site in order to activate the spike proteins for fusion with the host cell by achieving an irreversible conformational state. Furin-cleavage site makes SARS-CoV-2 S protein different from SARS-CoV S protein. This site comprises of four amino acid residues (P681, R682, R683, and A684) that is present at the boundary between the S1 and S2 subunit. Functionally, R682, R683, A684, and R685 create the nominal polybasic furin cleavage site (PRRAR). This sequence is not present in the previously known SARS-CoV and its related group 2b β-coronaviruses found in humans and this may be possibly one of the reasons for high virulence of SARS-CoV-2. Ubiquitously expressed furin proteases in humans are required for the activation of SARS-CoV-2 S protein. Genetically engineered furin-deleted SARS-CoV-2 spike protein has reduced processing and replication compared to the wild-type SARS-CoV-2 [11]. All investigators till date have shown that a stretch comprises of 14 amino acid residues (677QTNSPRRARSVASQ689) that includes PRRAR furin CS as well in Wt. SARS-CoV-2 S protein and its different variants found exist in form of a loop [20-22]

### SARS-CoV2 Omicron (B.1.1.529) variant

This newly emerged SARS-CoV2 variant has recently been found circulating in South Africa and Botswana. Omicron virus is unique from the previously characterized SARS-CoV2 virus. It has been found that it has 37 amino acid substitutions in the S protein and 15 substitutions among them are located in RBD and 10 among them in RBM, a part of RBD that interacts with ACE2 receptor. Compared to the original wild type SARS-CoV2, the dominant Delta (B.1.617.2) variant has only 2 mutations (L452R and T478K) in its RBM and K417N and E484K mutations occasionally. Therefore, Omicron variant may significantly impact the binding affinity to ACE2. Consequently, Omicron mutant has aroused wide concern; many countries have taken measures on entry restrictions to prevent its rapid spread. However, the transmissibility and immune evasion risk of Omicron have not been properly evaluated [23]. The spike protein of WT SARS-CoV-2 has 1273 amino acids (UniProt ID: P0DTC2), and its RBD is composed of residues 319–541 and RBM is of residues 437–507.2 The currently dominant Delta variant has only 4 mutations on its RBD (RBD Delta), much less than that on the Omicron RBD (RBD micron) (Fig. 1a). It could be seen that the 15 mutations of RBD Omicron are not evenly distributed in RBD, but rather crowed in its RBM with 10 residues, viz., N440K, G446S, S477N, T478K, E484A, Q493K, G496S, Q498R, N501Y and Y505H (Fig. 1a). By checking the effect of single mutation on ACE2 binding affinity reported by Bloom et al.3, it was found that 9 RBD Omicron mutations (S371L, S373P, S375F, K417N, G446S, E484A, G496S, Q498R, Y505H) should decrease the binding affinity to ACE2 while the other 6 mutations (G339D, N440K, S477N, T478K, Q493K, N501Y) should increase the binding affinity, resulting in a challenge of predicting its transmissibility and potential immune evasion risk [23, 24].

### α1AT protein

Alpha1 antitrypsin (α1AT) is a 52kDa serine-protease inhibitor, which abundantly synthesized by hepatocytes and to some extent by other cells of the human body. Neutrophils move towards the site of inflammation, release neutrophil elastase (protease). However, in case of α1AT deficiency, uncontrolled release of elastase leads to the manifestation of many diseases including parenchymal-degradation of lungs (emphysema).

Therefore, α1AT has an important role to neutralize the excessive production of elastase. X-ray crystallography, *insilico* analysis, and biochemical kinetics have provided us valuable insights to study its folding mechanism [25, 26]. During the initial folding process α1ATis kinetically trapped into a metastable state along with a patch of 15 amino acid residues (345-360) located near to its the C-terminus that forms a flexible loop called as reactive-center loop (RCL), which is exposed to polar solvent. The conformational plasticity of α1AT not only unveils its inhibitory activity but also provides a mechanistic understanding of pathogenic point-mutant α1AT susceptibility to misfolding and aggregation. The native fold of α1AT comprises of three β-sheets (A-C) enclosed by 8-9 α-helices (hA-hI), and RCL (connecting loop between β-s5A and β-s1C) is projecting out of the secondary structure of α1AT. The interaction of protease (neutrophil elastase) with the metastable α1AT leads to a drastic conformational transition upon protease-driven cleavage of RCL. The protease-driven cleavage of N-terminal portion at the P1□□P1 site in the RCL of the α1AT coupled with disabled protease by acylation steps into the β-sheet A (β-sA), thereby increasing the number of β-sheets from five to six. The deactivated protease bound with the thermodynamically stable α1AT is subsequently targeted for degradation [27].

### α1AT as a biomarker in SARS-CoV2 infected Patients

Reportedly, increased level of serum interleukin (IL)-6 cytokine has been observed in SARS-CoV2 infected patients who were observed to have moderate to severe symptoms. The IL-6/ α1AT ratio indicates a hemeostatic switch between pro- and anti-inflammatory responses. This ratio was seen prominently high in SARS-CoV2 infected patients admitted in the intensive care unit compared to stable ones. α1AT has been included as a potential clinical and biological biomarker marker under Clinical Trial number: NCT04348396 and NCT04366089 in Italy. Intriguingly, circulating levels of serum α1AT in SARS-CoV2-infected patients were significantly low compared to healthy ones. Interestingly, truncated α1AT mutant proteins were found to be significantly high in SARS-infected patients. The low bioavailablity of circulating α1AT in SARS-infected patients were found to have lung failure and can lead to acute respiratory distress syndrome [28].

### Potential role of α1AT in SARS-CoV2 infected patients

α1AT alleviates the acute lung damage by decreasing the IL-1β levels. It stands guard against the thrombotic complications such as macrothrombi and small vessel thrombosis observed in SARS-CoV2 infected patients. Thrombotic complications in SARS-CoV2-infected patients can lead to disease progression, organ failure, and poor outcome. It inhibits the cytokine storm and its augmentation therapy acts as an immune modulator to dampen the pro-inflammatory IL-6 cytokine levels and triggers balanced antiviral immune response that subsequently leads to the efficacious pathogen clearance without tissue damage. It sequesters IL-8 that consequently dampens the neutrophil influx and acute-lung damage and alleviates TGF-β-driven abrupt and longstanding damaging effects of SARS-CoV2. Accumulating investigations suggests conceivable association between α1AT and SARS-CoV2. FDA has approved α1AT as one of the drug that can be used with extraordinary clinical safety[29].

### TMPRSS2 (Transmembrane protease, serine 2)

TMPRSS2 is a 492 amino acid residue long single-pass type II membrane protein. It comprises of N-terminus domain located in the cytosol, TM domain, and a cytosolic domain. The cytosolic domain is further divided into three sub regions, namely LDL receptor class A from 113−148 amino acid that serves as a binding site for calcium, scavenger receptor cysteine-rich domain (from 149−242 amino acid residues), and a serine-protease (SP) domain extending from 255−492 amino acid residues of the S1 family.

The SP domain contains the H296, D354, and S441 that acts a catalytic triad. It is exclusively expressed at the cell surface and serves to regulate the cell-cell and cell matrix interaction. It is expressed in normal as well as in diseased condition. It is highly expressed in small intestine and to small extent in salivary gland, stomach, colon, and prostate gland. This protein is upregulated by androgenic hormones secreted by prostate cancer cells. It acts a receptor for specific ligands that helps it to coordinate with the extracellular environment of the cell. In lungs, TMPRSS2 has proposed to regulate the sodium current of the epithelial cells cleaving the sodium channel. It facilitates the SARS-CoV and SARS-CoV-2 infection in the host cells through two independent mechanisms, ACE2 receptor proteolytic cleavage that promotes viral infection, and cleavage of the SARS-CoV-2 S protein which activates the SARS-CoV-2 to get an access into the host cell.

### α1AT antiprotease inhibits the SARS-CoV2 infection by deactivating the TMPRSS2 protease activity

SARS-CoV-2 primarily spreads through inhalation of droplets and aerosols coming from the infected person and reinfection of respiratory tract cells with the same virus. In overwhelming majority of the SARS-CoV2 infected cases, infection remains confined to the upper airways causing no or mild symptoms. Chronic disease driven by SARS-CoV-2 is caused by the viral dissemination down to the lungs, consequently leading to the acute respiratory distress syndrome, cytokine burst, multi-organ failure, systemic septic shock, and death. The airway epithelium acts as a first-line of defence against the invading respiratory microorganisms through the mucociliary clearance action and its mucosal-associated lymphoid tissue. The epithelial lining fluid comprises of innate immunity effector molecules like short-peptides as well as proteins that act as antibacterial and antiviral agents, such as lactoferrin or defensins, lysozyme, and α1AT protein. Recently, it has been demonstrated that α1AT, a circulating serine protease inhibitor, stops the entry of SARS-CoV-2 in human airway epithelial by binding with TMPRSS2 antiprotease and deactivating it. Active TMPRSS2 activates the SARS-CoV-2 spike protein for its fusion with the human airway epithelial cells. *In vitro* experiments have revealed that α1AT protein inhibits the SARS-CoV-2 infection with IC_50_ of 10-20µM. Therefore, one would suggest that α1AT has anti-SARS-CoV-2 activity. It wouldn’t be surprising to suggest that α1AT deficient individuals are highly susceptible to SARS-CoV2 infection, more specifically, in early-onset COPD patients. Intravenous α1AT augmentation therapy has been used from decades in individuals suffering from α1AT deficiency driven early onset of parenchymal lung disease (emphysema). Therefore, α1AT augmentation therapy can prove instrumental by acting as an antinflamatory protein as well as anti-SARS-CoV2 agent. α1AT can be administered in body at significantly higher doses than those in routine α1AT deficiency. Interestingly, in some α1AT deficient patients, α1AT administered at doses of 120−250 mg/kg without causing any side effects, leads to a five-fold increase of α1AT concentration in lung epithelial lining fluid has been observed. However, it is still unclear whether α1AT infusion in human body will achieve a threshold concentration that is sufficient to block the entry of SARS-CoV-2 in lungs without leading to undesirable severe side effects remains yet to uncovered[28].

## *In silico* Methodology

### ACE2−SARS-CoV2-RBD interface analysis and electrostatic potential of various SARS-CoV2-RBD mutants

ACE2−SARS-CoV2-RBD (receptor binding domain) with the PBD: 6M0J, SARS-CoV2 trimeric spike protein closed (PDB: 6VXX) and open (PDB: 6VYB) conformation was analyzed through *PyMol* software.Various SARS-CoV2 mutants such as SARS-CoV2 (K417N), SARS-CoV2 (E484K), SARS-CoV2 (N501Y), SARS-CoV2 (E484Q), SARS-CoV2 (L452R), SARS-CoV2 double-mutant (E484Q and L452R), SARS-CoV2 triple-mutant (K417N, E484K, and N501Y) were introduced *in silico* through PyMol *software*. Electrostatic potential of wild-type SARS-CoV2 and its mutants were generated through PyMol *software*

### SARS-CoV2 Spike protein modeling

**P**rotein **H**omology/analogy **R**ecognition **E**ngine V 2.0 (*Phyre*^*2*^) was used for modeling the Original wild-type SARS-CoV2 S protein with PDB ID: 6vyb along with its subsequent mutation in B.1.1.7 at furin CS (HRRAR), B.1.617 (RRRAR) variants and MERS spike protein with the furin CS (PRSVR) was used as a template available structure for modeling. The modeled S protein with dimension (□): X=74.777, Y=124.039, Z=159.510 was built on template c7dk3B with 100% confidence and 83% coverage. 1062 residues (83% of the sequence) was modeled with 100% confidence by the single highest scoring template. 1141 residues (90%) was modeled at >90% confidence using multi-templates. SARS-CoV S protein with PDB ID: 5×58 was used as a template available structure for modeling. The modelled SARS-CoV S protein was found with dimension (□): X=80.71, Y=115.65, Z=148.49 was built on template c5×5bB with 100% confidence and 84% coverage. 1053 residues (84% of the sequence) was modeled with 100% confidence by the single highest scoring template. 1159 residues (92%) was modeled at >90% confidence using multi-templates [30].

### Protein□Protein docking

Single chain, water, ligand, and heme atom-free SARS-CoV2 spike protein (Modeled)−TMPRSS (PDB: 7MEQ; Cyan color) and α1AT (PDB ID:3NE4 (Green)−TMPRSS (Cyan) were prepared through the *<u>Discovery Studio Visualizer</u>. <u>Chimera software</u>* was used to add charge to the modelled SARS-CoV2 single chain spike protein, TMPRSS, furin and α1AT protein.

The prepared proteins were docked through *https://cluspro.bu.edu*. [31-33]The energies of various protein-protein docked complex were as follows: electrostatic-favored energy (TMPRSS2−SARS CoV2 spike protein) =0.40E_rep_+−0.40E_att_+1200E_elec_+1.00E_DARS_, VdW+Elec (TMPRSS2−SARS-CoV2) =0.40E_rep_+ −0.10E_att_+600E_elec_+0.00E_DARS_, electrostatic favored (Furin−SARS-CoV2 spike protein)=0.40Erep+−0.40E_att_+1200E_elec_+1.00E_DARS_, and electrostatic-favored (Furin−α1AT) =0.40E_rep_+ −0.40E_att_+1200E_elec_+1.00E_DARS_. The docked complexes were analyzed for hydrogen bonding between the protein interfaces through PyMol software [34].

### SARS-CoV2 Spike protein Arg/Lys cleavage prediction at S1/S2 furin cleavage site

Propeptide cleavage sites predicted (ProP): Arg(R)/Lys (K): 3. If the score is >0.5, the residue is predicted to be followed by propeptide cleavage site.The higher the score the more confident the prediction. Prediction scores for the S1/S2 furin cleavage site in S protein were analyzed by using the *<u>ProP 1.0 server</u>* (*www.cbs.dtu.dk/services/ProP/*)[35]

## Results

### Unique electrostatic potential of SARS-CoV2 variants compared to its original wild-type

We tried to observe the electrostatic potential of SARS-CoV2 RBD mutants complexed with the ACE2 receptor at their binding interface. We observed a dramatic change in the electrostatic potential of SARS-CoV2 RBD (K417N) and SARS-CoV2 RBD triple mutant (K417N, E484K, N501Y) complexed with the ACE2 receptor at their binding interface compared to the wild-type SARS-CoV2 RBD (**Fig 2A-H**). Our results revealed that single and triple-mutants of SARS-CoV2 RBD extensively differ in electrostatic potential at their binding interface.

**Fig 2.**
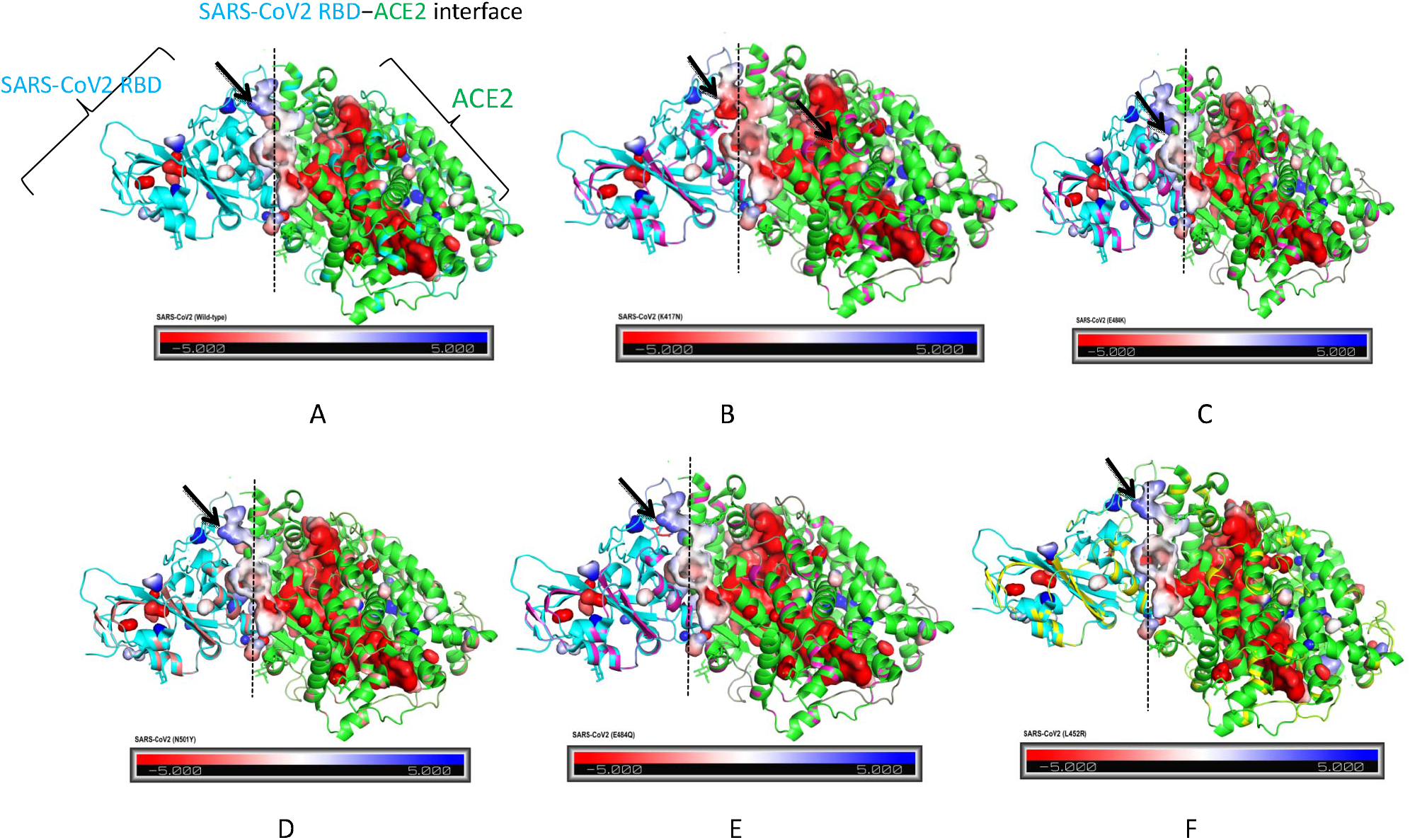
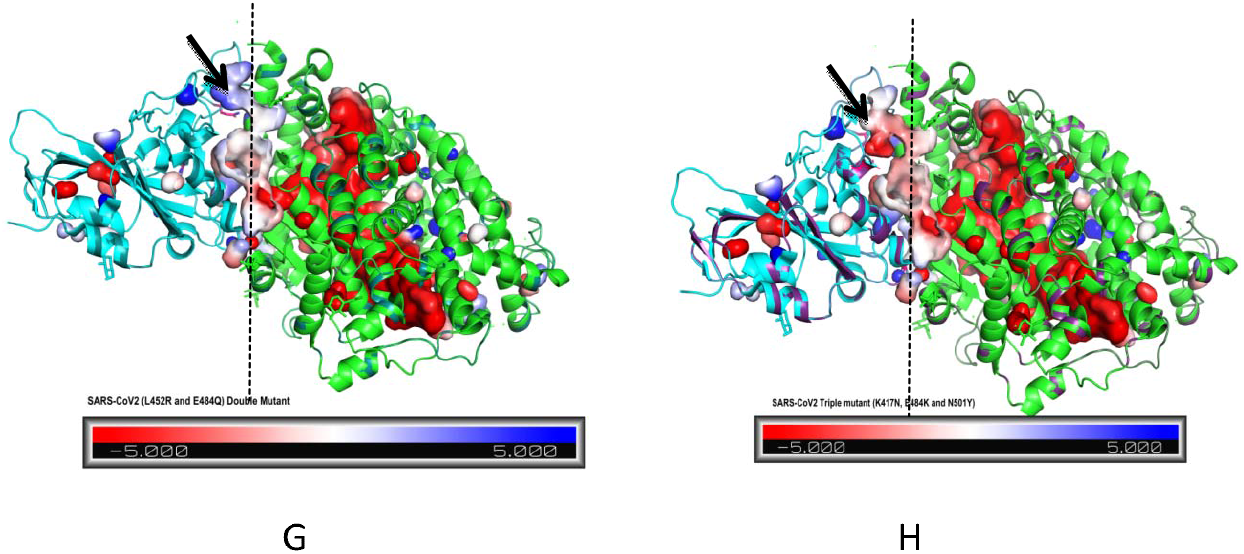
Electrostatic-potential of wild-type SARS-CoV2 and its variants. (**A**) Electrostatic potential of wild-type SARS-CoV2. (**B−H**) Electrostatic potential of SARS-CoV2 (K417N), SARS-CoV2 (E484K), SARS-CoV2 (N501Y), SARS-CoV2 (E484Q), and SARS-CoV2 (L452R). (**G**) SARS-CoV2 double-mutant (L452R and E484Q). (**H**) SARS-CoV2 triple-mutant (K417N, E484K and N501Y). (**I**) Electrostatic potential of wild-type SARS-CoV2 without K417N mutation.

### 14 amino acid missing residues form antiparallel β-sheet harboring furin cleavage site

Phylogenetic investigation of the newly evolved SARS-CoV-2 is unique in a manner that it has an insertion of RRAR furin cleavage site at the S1/S2 site of SARS-CoV2 S protein compared to previously identified SARS-CoV and other SARS-related coronaviruses. The addition of this sequence in SARS-CoV2 has been speculated to provide a gain-of-function, thereby increasing its efficacy of transmission in humans by easily gaining an access to airway epithelial cells compared to other lineage that eventually leads to a host of clinical condition such as lung damage, bilobular-pneumonia, macrothrombi formation etc. N-terminal domain and RBD harboring RB motiff are present in the S1 subunit of SARS-CoV-2 S protein. The S2 subunit of SARS-CoV-2 consists of fusion peptide, heptad repeat 1, central helix, connector domain, heptad repeat 2, transmembrane domain, and a tail flanking towards the cytosol (**Fig.3A**). The furin cleavage site resides between these two domains. In overwhelming majority of the coronavirus, host proteases like TMPRSS2 cleave the furin CS through an irreversible conformational change that assists in the viral fusion with the host cell. Majority of investigators have reported the presence of a loop between S1/S2 junction of SARS-CoV2 S protein. In this connection, we modeled full SARS-CoV2 spike protein through Protein Homology/ analogy Y Recognition Engine V 2.0) by taking the available SARS-CoV2 spike protein with PDB: 6VYB as a template. SARS-CoV2 S protein was modeled because of the reason that the available structure lacks a nine amino acid patch (677<u>QTNSPRRAR</u>685) that essentially provides a furin cleavage site. We found that this sequence forms two anti-parallel β-sheets comprising of FCS that protrudes out of the main body of SARS-CoV2 S protein (**Fig 3C**). The electrostatic potential of the available and modeled SARS-CoV2 S protein is shown in **Fig3. B, D**. The secondary structure of our modeled SARS-CoV2 S protein has 37% in β strand, 26% in α helices, 16% in a disordered state, and 7% existing as transmembrane (**Fig 3E**.). We anticipate that this β-sheet is used as a scaffold by proteases to act on furin-CS in SARS-CoV2 S protein.

**Fig 3.**
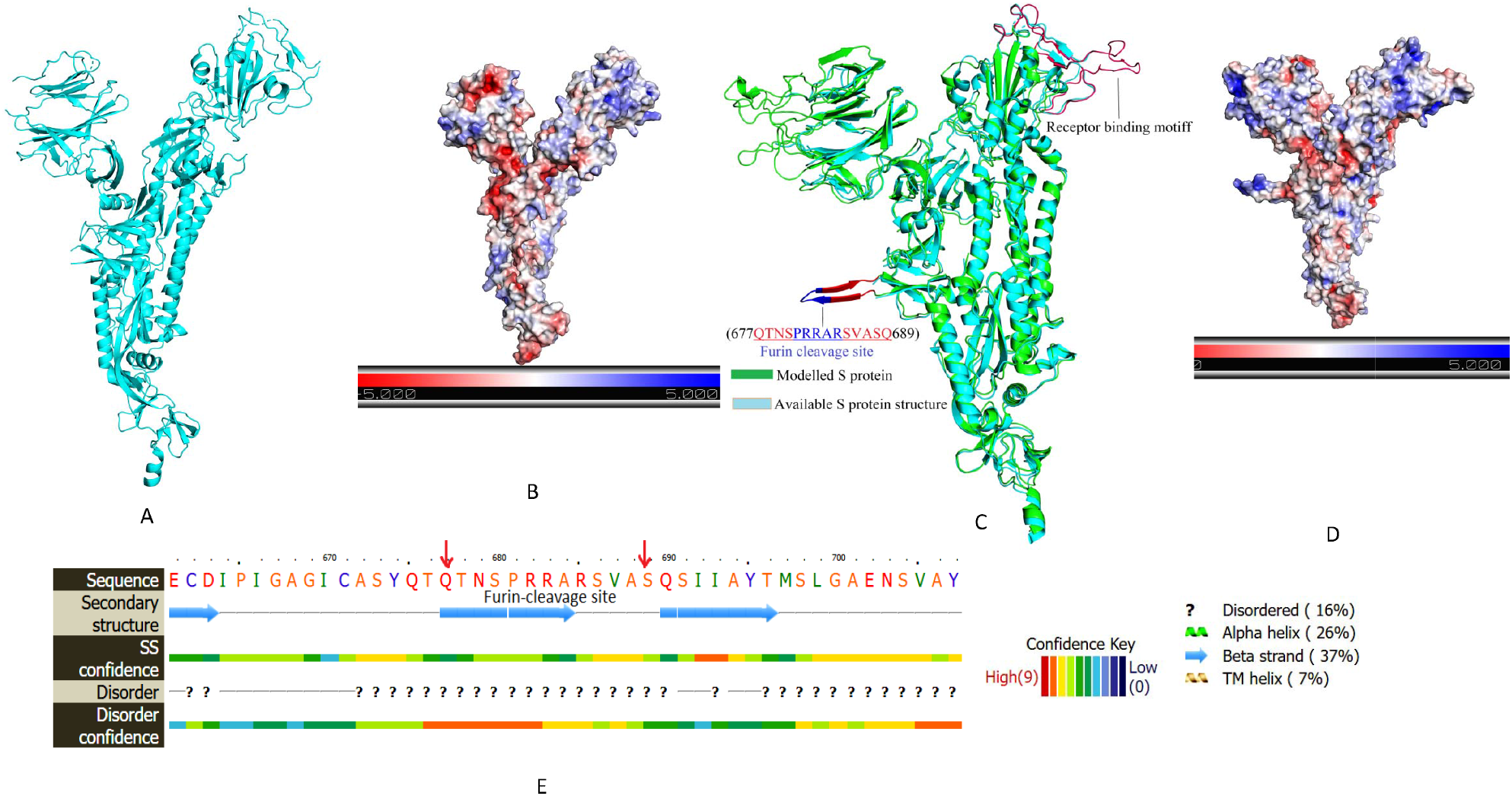
Relative comparison of the SARS-CoV2 Spike protein against the modeled one showing furin-cleavage site harboring in an anti parallel β-sheet. (**A**) Structure of the available SARS-CoV2 Spike protein with PDB ID:6vyb along with its electrostatic potential (**B**). (**C**) Structure of the modelled SARS-CoV2 Spike protein (Green) aligned with the the available SARS-CoV2 Spike protein (Cyan). Anti parallel β-sheet (highlighted with Red color) encompasses furin-cleavage site 677<u>QTNSPRRAR</u>685 (highlighted with Blue color). (**D**) Electrostatic potential of the modeled protein. (**E**) SARS-CoV2 Spike protein sequence showing a β-stand housing furin-cleavage site along with the overall percentage of secondary structure in this protein.

### Relative comparison of the SARS-CoV, MERS, SARS-CoV2 and its furin CS variants differs in percentage secondary structure

We modeled the SARS-CoV, MERS, SARS-CoV2 and its furin CS variants (HRRAR and RRRAR). We found that MERS S protein also harbors antiparallel β-sheet whose percentage secondary structure resembled closely with the SARS-CoV2 spike protein with Furin CS HRRAR B.1.1.7 variant (**Fig** 4. **G, I, L, N**). While switching over from Wt. SARS-CoV2 with PRRAR furin CS to B.1.1.7 variant with HRRAR furin CS, we found 3% increase in the percentage content of β stands. We found a visible difference in the percentage secondary structure between SARS-CoV and SARS-CoV2 (**Fig 4. F, H, K, M**). Figure **J, O** shows the percentage secondary structure of SARS-CoV2 HRRAR B.1.1.7 variant. The superimposed spike protein of SARS-CoV, MERS, SARS-CoV2 and its furin CS variants (HRRAR and RRRAR) is gown in Figure 4**A**. A close overview of supermposed β sheets is shown in Figure **4B**.

**Fig 4.**
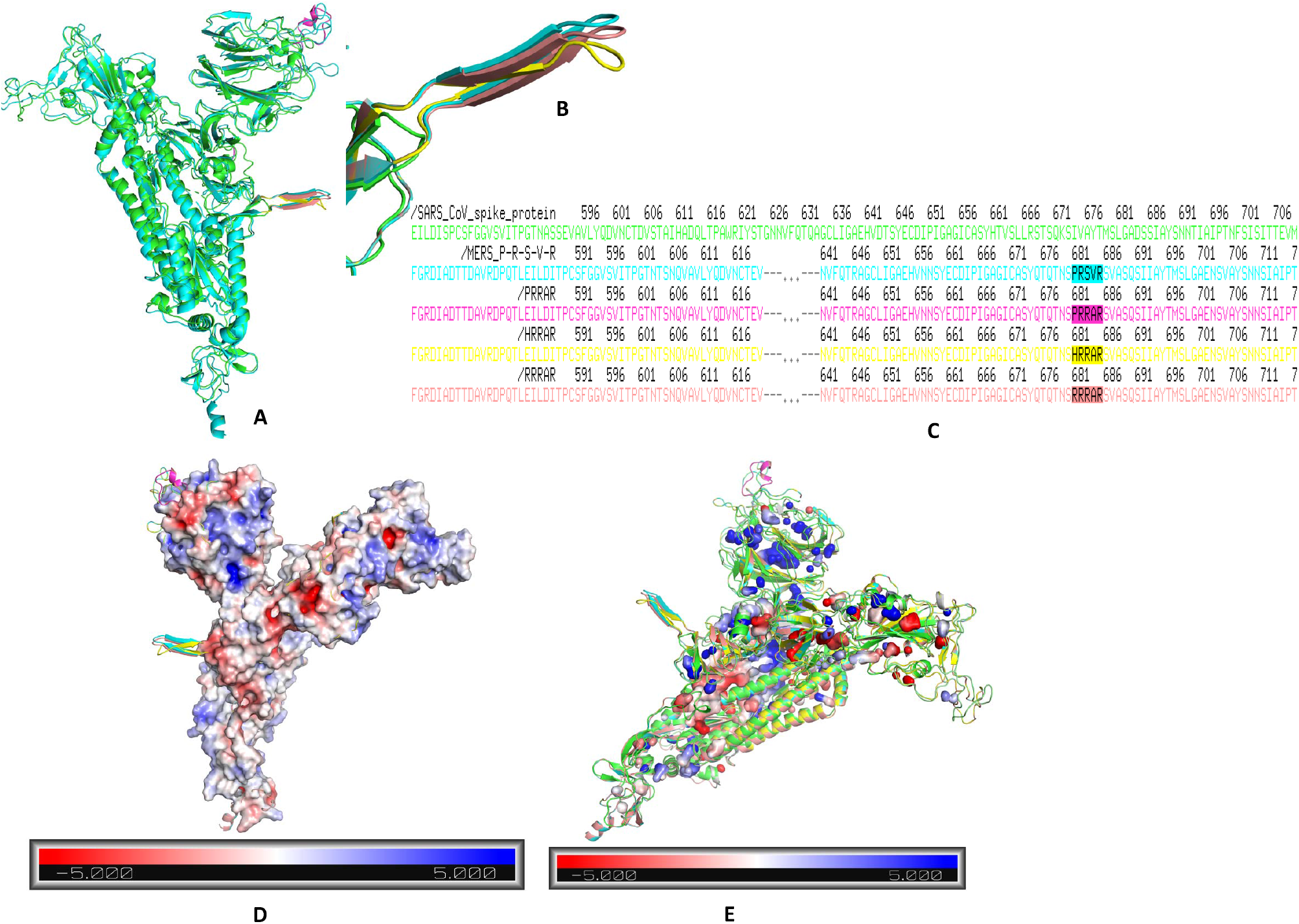
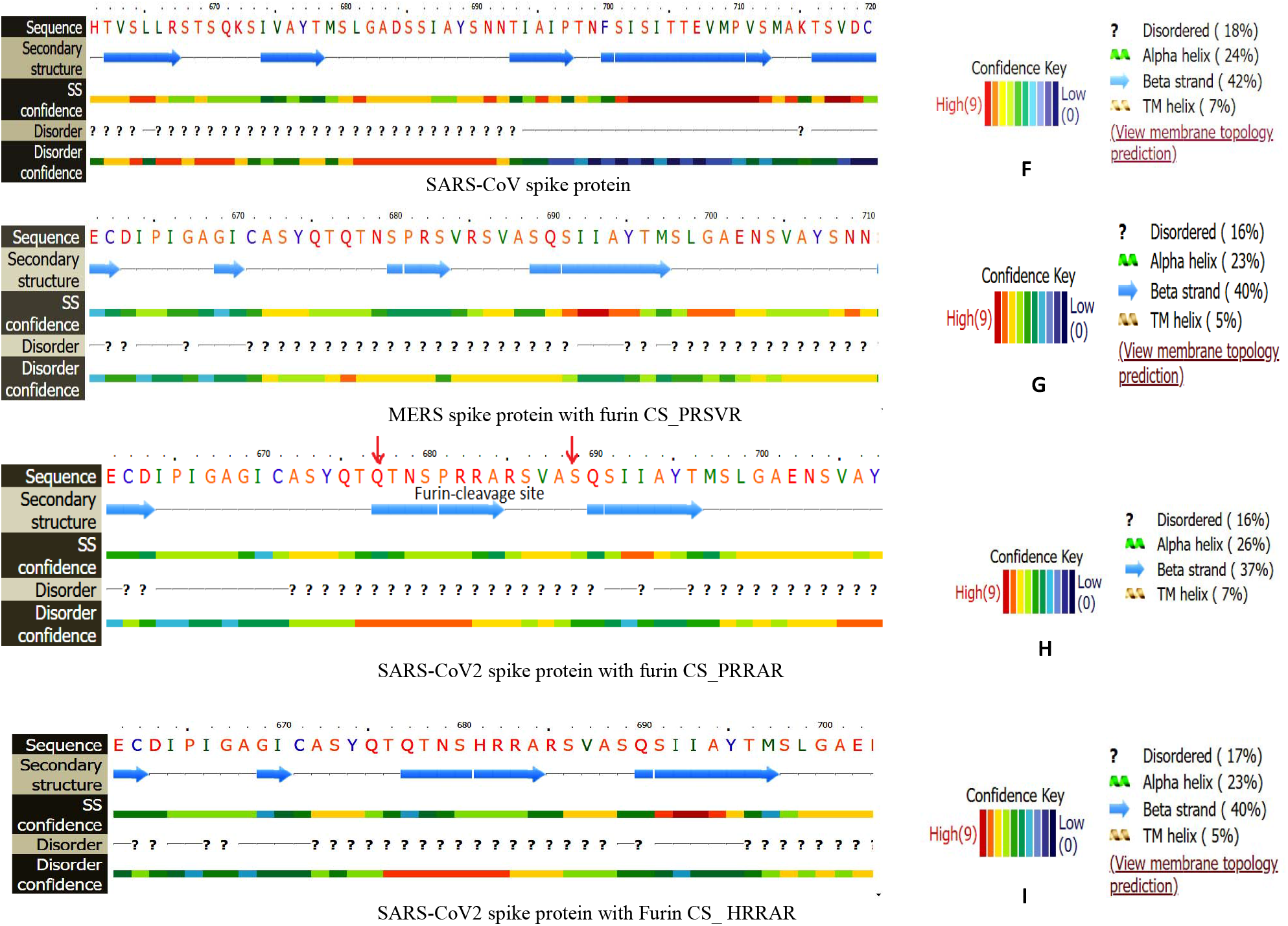
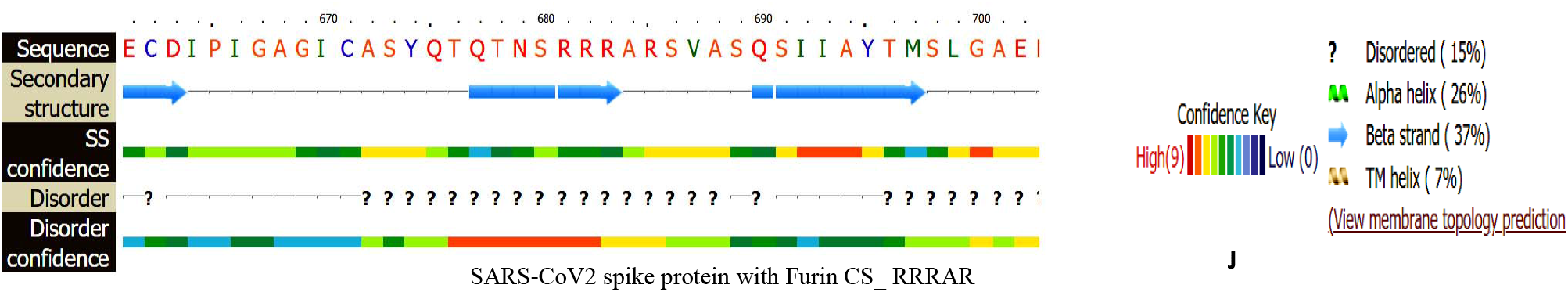
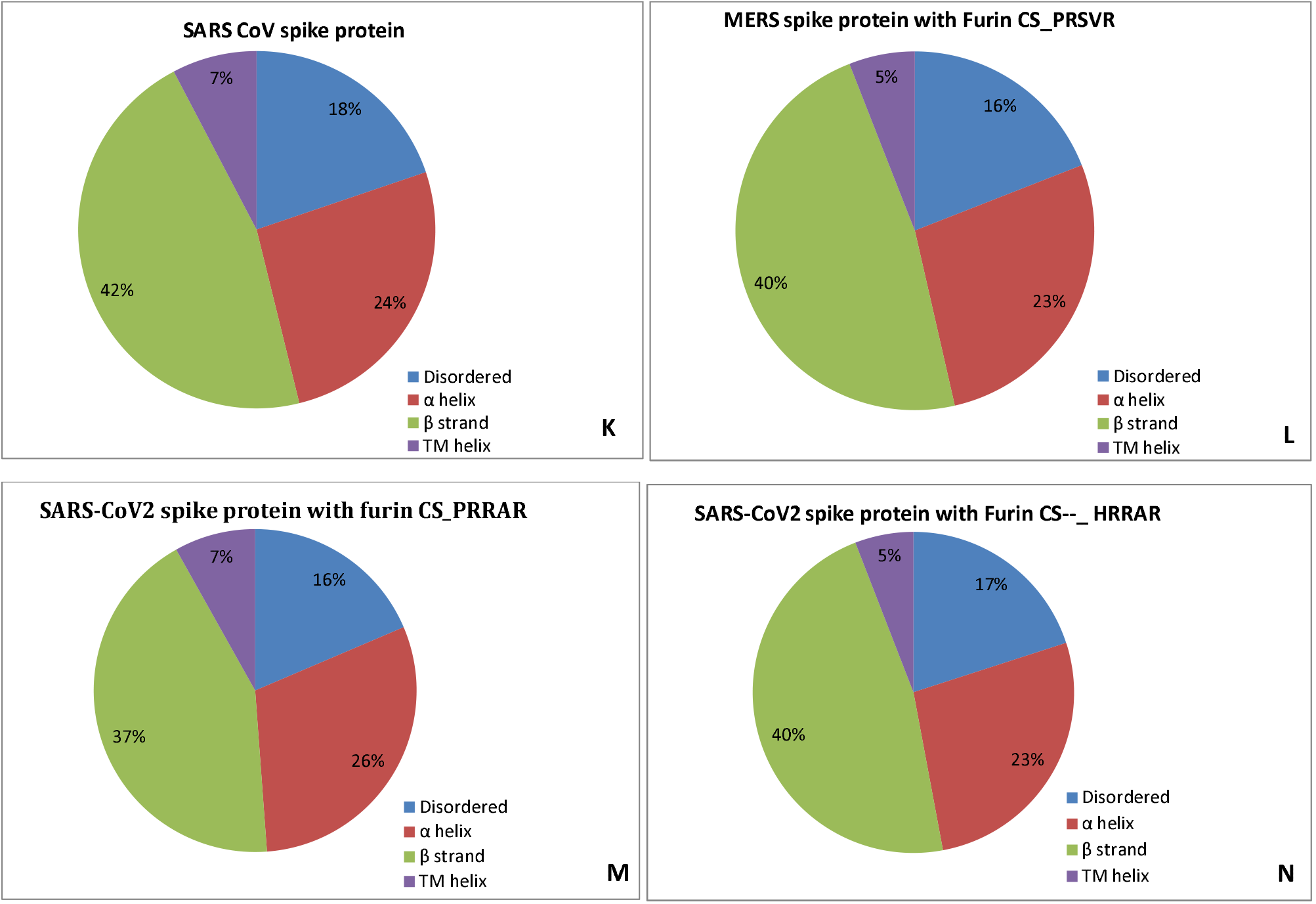
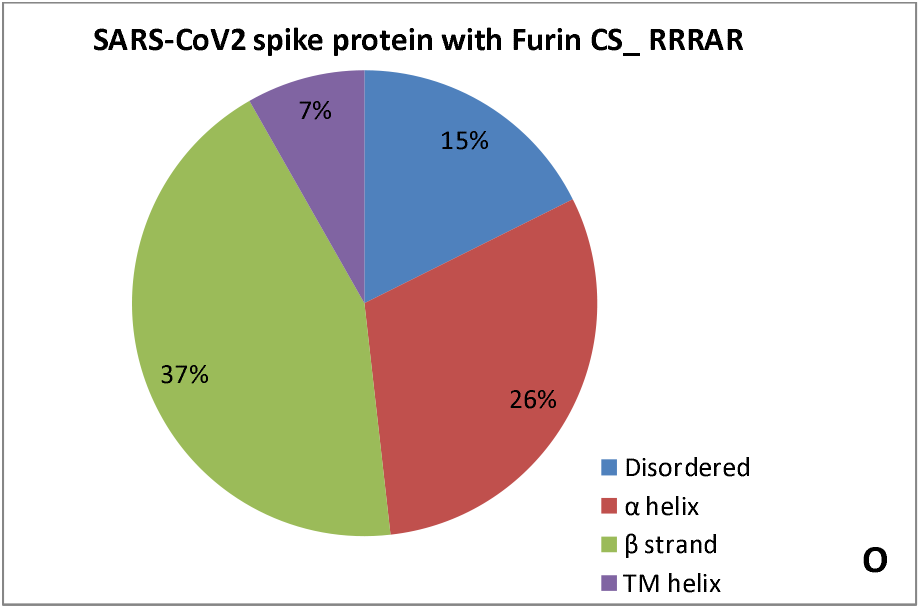
Relative comparison of the SARS-CoV spike protein, MERS spike protein, SARS-CoV2 spike protein with different furin cleavage sites (**A**) Superimposed SARS-CoV spike protein, MERS spike protein with furin CS (PRSVR), SARS-CoV2 spike protein with different furin cleavage (PRRAR, HRRAR, RRAR) (**B**) High-resolution image of superimposed β stands. (**C**)Amino acid sequence alignment of the SARS-CoV spike protein, MERS spike protein, SARS-CoV2 spike protein with different furin cleavage sites highlighted with their respective colors. (**D, E**) Electrostatic potential of superimposed MERS spike protein with furin CS (PRSVR), SARS-CoV2 spike protein with different furin cleavage (PRRAR, HRRAR, RRAR). (**F-J**) Secondary structure features of SARS-CoV spike protein, MERS spike protein, SARS-CoV2 spike protein with different furin cleavage sites. (**K-O**) Graphical representation of SARS-CoV spike protein, MERS spike protein, SARS-CoV2 spike protein with different furin cleavage sites highlighting their percentage secondary structure variations. TM-transmembrane

The amino acid sequence alignment of SARS-CoV, MERS, SARS-CoV2 and its furin CS variants (HRRAR and RRRAR) is gown in Figure **4C**. Electrostatic potential of superimposed SARS-CoV, MERS, SARS-CoV2 and its furin CS variants (HRRAR and RRRAR) spike protein is shown in Figure **4 D, E**. Interestingly, we found that the change of B.1.1.7 to B.1.617 variant comprising of RRRAR furin CS shifted the percentage secondary structure back to that found in Wt. SARS-CoV2. Overall, we infer that the recently emerged SARS-CoV2 variant like B.1.617 RRRAR variant is not showing any dramatic change and possess the similar percentage secondary structure as found original Wt-type SARS-CoV2 S protein

### RRAR furin sequence of SARS-CoV2 S protein interacts with the TMPRSS2 protease

We found that SARS-CoV2 S protein comprising of furin cleavage site (682<u>RRAR685</u>) interacts with the D417, N418, Y416, and N418 amino acid residues of the TMPRSS2 protease (**Fig 5D, E**). The electrostatic surface potential of the SARS-CoV2 S protein□TMPRSS2 protease is shown in **Fig 5F**.

**Fig 5.**
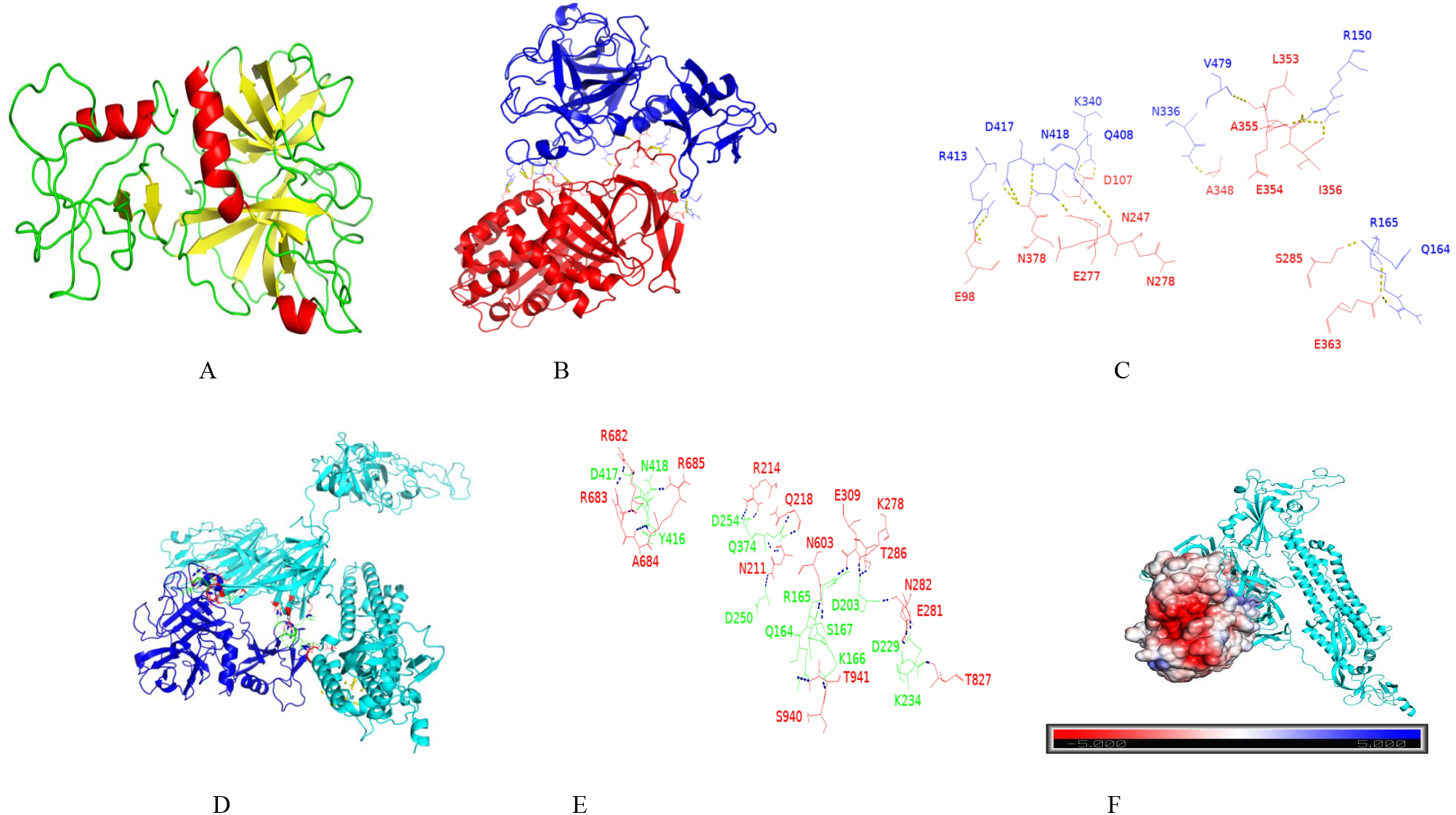
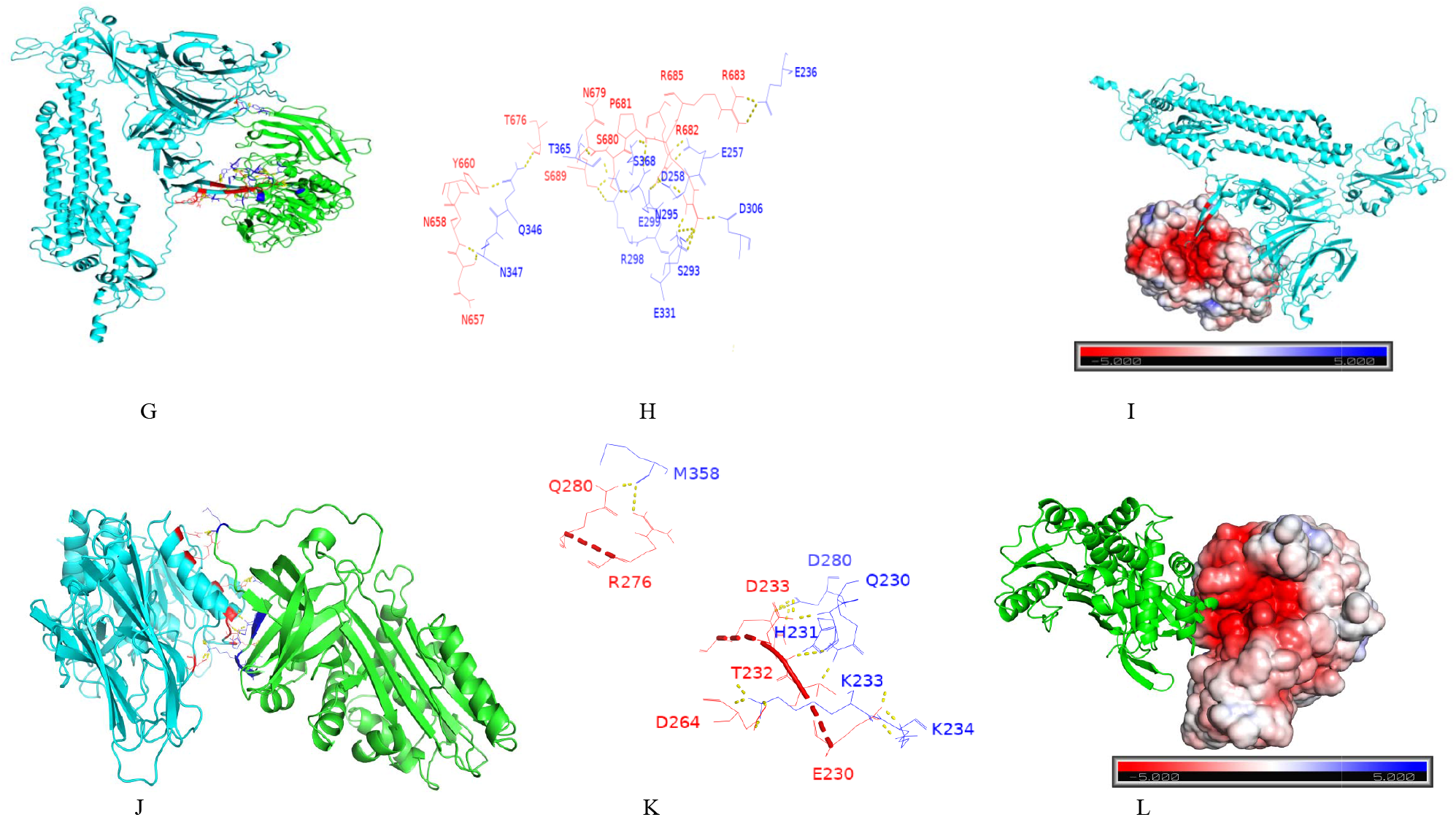
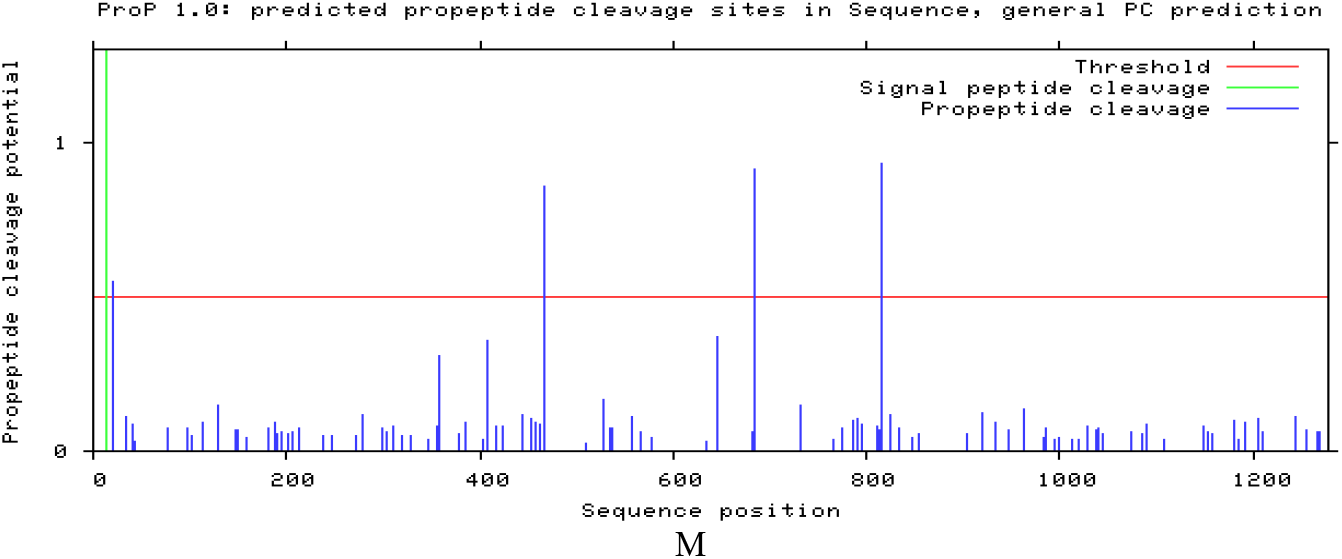
(**A**) Single chain TMPRSS2 (**PDB: 7MEQ**) prepared through Maestro-Schrodinger software (trail-license provided by www.schrodinger.com/products/maestro to Arif Bashir) for addition of charge and missing loops. α1AT (**PBD ID: 3NE4**) was also prepared through protein-prep wizard, Maestro Schrodinger. Protein-protein docking performed on supercomputer. The access to the supercomputer was provided by www.cluspro.bu.edu (**B**) TMPRSS2−α1AT docked molecule. (**C**) TMPRSS2−α1AT interacting amino acid residues. Docking was performed through https://cluspro.bu.edu. Electrostatic-favored=E=0.40E_rep_+ −0.40E_att_+1200E_elec_+1.00E_DARS_. (**D**) TMPRSS2 (Blue)−SARS-CoV2 spike protein (Cyan) docked molecule. (**E**) TMPRSS2− SARS-CoV2 spike protein interacting amino acid residues. VdW+Elec= E=0.40E_rep_+ −0.10E_att_+600E_elec_+0.00E_DARS_. (**F**) Electrostatic potential of TMPRSS2− SARS-CoV2 spike protein (Cyan). (**G**) Furin (Green)−SARS-CoV2 (Cyan) spike protein(Cyan) docked molecule. Electorstatic favored=0.40Erep+ −0.40E_att_+1200E_elec_+1.00E_DARS_. (**H**) Furin (Green)−SARS-CoV2 (Red) spike protein (Blue) interacting amino acids. (**I**) Electrostatic potential of Furin−SARS-CoV2 spike protein (Cyan). (**J**) Furin (Cyan)− α1AT (Green) spike docked protein. Electrostatic-favored=0.40E_rep_+ −0.40E_att_+1200E_elec_+1.00E_DARS_ (**K**) Furin (Red) and α1AT (Blue) interacting amino acids. (**L**) Electrostatic potential of Furin−α1AT (Green). (**M**) Graph showing the SARS-CoV2 Spike protein Arg/Lys cleavage prediction at S1/S2 furin cleavage site. Propeptide cleavage sites predicted Proline: Arginine/Lysine: 3. If the score is >0.5, the residue is predicted to be followed by propeptide cleavage site. The higher the score the more confident the prediction. Prediction scores for the S1/S2 furin cleavage site in S protein were analyzed by using the *<u>ProP 1.0 server</u>* (www.cbs.dtu.dk/services/ProP/).

### PRRAR furin sequence of SARS-CoV2 S protein interacts with the furin protease

We found that SARS-CoV2 S protein comprising of furin cleavage site (681P<u>RR-R685</u>) interacts with the S368, D258, E257, and E236 amino acid residues of the furin protease (**Fig 5G, H**). The electrostatic surface potential of the SARS-CoV2 S protein□furin protease is shown in **Fig 5I**.

### Arg/Lys cleavage prediction at S1/S2 furin cleavage site in SARS-CoV2 S protein

We analysed the Arginine (Arg)/Lysine (Lys) cleavage prediction at S1/S2 furin cleavage site in SARS-CoV2 S protein. The propeptide cleavage sites predicted Proline: Arg/Lys ratio 3, pointing the three prominent Arg/Lys cleavage sites in SARS-CoV2 S protein at around between 400-600, 600-800, and 800-820 amino acid residues (**Fig 5M**).

### Reactive-center loop of α1AT interacts with TMPRSS2 attached on the surface of airway epithelial cells

The C-terminal 15 amino acid residues forms a reactive center loop (RCL) in α1AT. This loop holds a key position in α1AT by interacting with the neutrophil elastase (NE), a protease that is released at the site of inflammation. In addition to NE, α1AT interacts with a wide range of proteases including the TMPRSS2 that is located on the surface of the airway epithelial cells. The interaction of proteases with the α1AT cleaves the RCL, abrogates the α1AT antiprotease activity and inserts this loop between the α-helices, thereby increasing the number of β-sheets of the α1AT from five to six. In this context, α1AT acts as an antiprotease effector protein to dampen the immune response driven by the activation of NE during microbial invasion that leads to inflammation. TMPRSS2 binds with the SARS-CoV2 S proteins S1 subunit bound with the ACE receptor present on airway epithelial cells. The bound TMPRSS2 cleaves the S protein at S1 site. We found that α1AT by virtue of its RCL interacts with the TMPRSS2 and deactivates its protease activity (**Fig 5A, B, C**).

### Reactive-center loop of α1AT antiprotease interacts with furin protease

We observed that α1AT uses its RCL to interact and subsequently deactivate the furin protease (**Fig 5J, J**). The electrostatic potential of the complex is shown in **Fig 5L**.

## Discussion

Majority of investigators have reported the presence of a loop harboring the furin cleavage site at S1/S2 junction of SARS-CoV2 S protein. We are for the first time reporting this patch comprises of 14 amino acid residues (677QTNSPRRARSVASQ689) that form an antiparallel β-sheet comprising of already known PRRAR furin polybasic CS. Our results revealed that single and triple-mutants of SARS-CoV2 RBD extensively differ in electrostatic potential at their binding interface. We found that MERS S protein also harbors antiparallel β-sheet whose percentage secondary structure resembled closely with the SARS-CoV2 spike protein with Furin CS HRRAR B.1.1.7 variant. We observe that the recently emerged SARS-CoV2 variant like B.1.617 RRRAR variant is not showing any dramatic change and possess the similar percentage secondary structure as found original Wt-type SARS-CoV2 S protein. We anticipate that the anti-parallel β-sheet formed by SARS-CoV, MERS, SARS-CoV2 and its furin CS variants (HRRAR and RRRAR) spike protein is used as a scaffold by proteases to act on furin-CS in SARS-CoV2 S protein. Additionally, we studied the interaction of modeled SARS-CoV2 S protein with transmembrane protease, serine 2 (TMPRSS2), and furin proteases, which clearly advocated that proteases like furin and TMPRSS2 cleaves furin CS sequence (PRRAR) located in antiparallel β-sheet of our modeled SARS-CoV2 S protein at its S1/S2 junction.

## Contributions

The idea and proposal of the study was designed by Arif Bashir

## Acknowledgments

None

## References

1. Zhou, P., et al., A pneumonia outbreak associated with a new coronavirus of probable bat origin. Nature. 2020 Mar;579(7798):270–273. doi(2020 Feb 3): p. 10.1038/s41586-020-2012-7.

2. Wu, F., et al., A new coronavirus associated with human respiratory disease in China. Nature. 2020 Mar;579(7798):265–269. doi(2020 Feb 3): p. 10.1038/s41586-020-2008-3.

3. Zhu, N., et al., A Novel Coronavirus from Patients with Pneumonia in China, 2019. N Engl J Med. 2020 Feb 20;382(8):727–733. doi(2020 Jan 24): p. 10.1056/NEJMoa2001017.

4. Lu, R., et al., Genomic characterisation and epidemiology of 2019 novel coronavirus: implications for virus origins and receptor binding. Lancet. 2020 Feb 22;395(10224):565–574. doi(2020 Jan 30): p. 10.1016/S0140-6736(20)30251-8.

5. Harvey, W.T., et al., SARS-CoV-2 variants, spike mutations and immune escape. Nat Rev Microbiol. 2021 Jul;19(7):409–424. doi(2021 Jun 1): p. 10.1038/s41579-021-00573-0.

6. Bashor, L., et al., SARS-CoV-2 evolution in animals suggests mechanisms for rapid variant selection. Proc Natl Acad Sci U S A, (10): p. 2021 Nov 2;118(44):e2105253118.

7. Sun, F., et al., SARS-CoV-2 Quasispecies Provides an Advantage Mutation Pool for the Epidemic Variants. Microbiol Spectr. 2021 Sep 3;9(1):e0026121. doi(2021 Aug 4): p. 10.1128/Spectrum.00261-21.

8. Gui, M., et al., Cryo-electron microscopy structures of the SARS-CoV spike glycoprotein reveal a prerequisite conformational state for receptor binding. Cell Res. 2017 Jan;27(1):119–129. doi(2016 Dec 23): p. 10.1038/cr.2016.152.

9. Kirchdoerfer, R.N., et al., Stabilized coronavirus spikes are resistant to conformational changes induced by receptor recognition or proteolysis. Sci Rep, (10): p. 2018 Oct 24;8(1):15701.

10. Lan, J., et al., Structure of the SARS-CoV-2 spike receptor-binding domain bound to the ACE2 receptor. Nature. 2020 May;581(7807):215–220. doi(2020 Mar 30): p. 10.1038/s41586-020-2180-5.

11. Peacock, T.P., et al., The furin cleavage site in the SARS-CoV-2 spike protein is required for transmission in ferrets. Nat Microbiol. 2021 Jul;6(7):899–909. doi(2021 Apr 27): p. 10.1038/s41564-021-00908-w.

12. Yuan, Y., et al., Cryo-EM structures of MERS-CoV and SARS-CoV spike glycoproteins reveal the dynamic receptor binding domains. Nat Commun, (10): p. 2017 Apr 10;8:15092.

13. Walls, A.C., et al., Structure, Function, and Antigenicity of the SARS-CoV-2 Spike Glycoprotein. Cell, (10): p. 2020 Dec 10;183(6):1735.

14. Wrapp, D., et al., Cryo-EM structure of the 2019-nCoV spike in the prefusion conformation. Science. 2020 Mar 13;367(6483):1260–1263. doi(2020 Feb 19): p. 10.1126/science.abb2507.

15. Wan, Y., et al., Receptor Recognition by the Novel Coronavirus from Wuhan: an Analysis Based on Decade-Long Structural Studies of SARS Coronavirus. J Virol. 2020 Mar 17;94(7):e00127–20. doi(2020 Mar 17): p. 10.1128/JVI.00127-20.

16. Andersen, K.G., et al., The proximal origin of SARS-CoV-2. Nat Med, (10): p. 2020 Apr;26(4):450–452.

17. Madu, I.G., et al., Characterization of a highly conserved domain within the severe acute respiratory syndrome coronavirus spike protein S2 domain with characteristics of a viral fusion peptide. J Virol. 2009 Aug;83(15):7411–21. doi(2009 May 13): p. 10.1128/JVI.00079-09.

18. Whittaker, G.R., SARS-CoV-2 spike and its adaptable furin cleavage site. Lancet Microbe. 2021 Oct;2(10):e488–e489. doi(2021 Aug 6): p. 10.1016/S2666-5247(21)00174-9.

19. Hulswit, R.J., C.A. de Haan, and B.J. Bosch, Coronavirus Spike Protein and Tropism Changes. Adv Virus Res. 2016;96:29–57. doi(2016 Sep 13): p. 10.1016/bs.aivir.2016.08.004.

20. Johnson, B.A., et al., Furin Cleavage Site Is Key to SARS-CoV-2 Pathogenesis. bioRxiv. 2020 Aug 26:2020.08.26.268854. doi: p. 10.1101/2020.08.26.268854.

21. Johnson, B.A., et al., Loss of furin cleavage site attenuates SARS-CoV-2 pathogenesis. Nature. 2021 Mar;591(7849):293–299. doi(2021 Jan 25): p. 10.1038/s41586-021-03237-4.

22. Wu, Y. and S. Zhao, Furin cleavage sites naturally occur in coronaviruses. Stem Cell Res, (10): p. 2020 Dec 9;50:102115.

23. Miller, N.L., et al., Insights on the mutational landscape of the SARS-CoV-2 Omicron variant. bioRxiv. 2021 Dec 10:2021.12.06.471499. doi: p. 10.1101/2021.12.06.471499.

24. Wu, L., et al., SARS-CoV-2 Omicron RBD shows weaker binding affinity than the currently dominant Delta variant to human ACE2. Signal Transduct Target Ther, (10): p. 2022 Jan 5;7(1):8.

25. Krishnan, B. and L.M. Gierasch, Dynamic local unfolding in the serpin alpha-1 antitrypsin provides a mechanism for loop insertion and polymerization. Nat Struct Mol Biol. 2011 Feb;18(2):222–6. doi(2011 Jan 23): p. 10.1038/nsmb.1976.

26. Tsutsui, Y., R. Dela Cruz, and P.L. Wintrode, Folding mechanism of the metastable serpin alpha1-antitrypsin. Proc Natl Acad Sci U S A. 2012 Mar 20;109(12):4467–72. doi(2012 Mar 5): p. 10.1073/pnas.1109125109.

27. Bashir, A., et al., Aggregation of M3 (E376D) variant of alpha1-antitrypsin. Sci Rep, (10): p. 2020 May 19;10(1):8290.

28. Wettstein, L., et al., Alpha-1 antitrypsin inhibits TMPRSS2 protease activity and SARS-CoV-2 infection. Nat Commun, (10): p. 2021 Mar 19;12(1):1726.

29. Azouz, N.P., et al., Alpha 1 Antitrypsin is an Inhibitor of the SARS-CoV-2-Priming Protease TMPRSS2. Pathog Immun. 2021 Apr 26;6(1):55–74. doi(2021): p. 10.20411/pai.v6i1.408.

30. Kelley, L.A., et al., The Phyre2 web portal for protein modeling, prediction and analysis. Nat Protoc. 2015 Jun;10(6):845–58. doi(2015 May 7): p. 10.1038/nprot.2015.053.

31. Desta, I.T., et al., Performance and Its Limits in Rigid Body Protein-Protein Docking. Structure. 2020 Sep 1;28(9):1071-1081.e3. doi(2020 Jul 9): p. 10.1016/j.str.2020.06.006.

32. Vajda, S., et al., New additions to the ClusPro server motivated by CAPRI. Proteins. 2017 Mar;85(3):435–444. doi(2017 Jan 5): p. 10.1002/prot.25219.

33. Kozakov, D., et al., The ClusPro web server for protein-protein docking. Nat Protoc. 2017 Feb;12(2):255–278. doi(2017 Jan 12): p. 10.1038/nprot.2016.169.

34. Seeliger, D. and B.L. de Groot, Ligand docking and binding site analysis with PyMOL and Autodock/Vina. J Comput Aided Mol Des. 2010 May;24(5):417–22. doi(2010 Apr 17): p. 10.1007/s10822-010-9352-6.

35. Duckert, P., S. Brunak, and N. Blom, Prediction of proprotein convertase cleavage sites. Protein Eng Des Sel, (10): p. 2004 Jan;17(1):107–12.

